# Spatial distance and temporal attentional focus modulate voluntary action preparation and awareness

**DOI:** 10.1101/2025.05.20.654810

**Authors:** Gaiqing Kong, Bastien Barlerin, Clément Desoche, Luke Miller, Francesco Pavani, Alessandro Farnè, Marine Vernet

## Abstract

Peripersonal space (PPS)—the immediate space surrounding the body—modulates perception and motor control. However, its impact on how voluntary actions are initiated and subjectively experienced remains underexplored. Similarly, how directing attention to different phases of an action, such as intention formation versus execution, and anticipating outcomes of an action, modulates the neural readiness for movement, has yet to be fully examined. This study investigates whether spatial proximity, attentional focus, and anticipated action outcomes influence action preparation and action awareness, using a virtual reality adaptation of the Libet clock paradigm during EEG recordings. Neural results reveal that attentional focus and anticipated action outcomes modulate different phases of motor preparation, as indexed by the readiness potential (RP)—a gradual buildup of neural activity preceding voluntary movement. Focusing on decision timing (without subsequent action outcomes) enhances early RP amplitude and decreases the late RP slope, suggesting increased preparatory neural engagement during intention formation. In contrast, focusing on action execution leads to a steeper late RP slope, indicating later and faster motor activity buildup when attention is directed toward movement onset. Anticipating action outcomes increased late RP slope, which was accompanied by the temporal binding effect: when a tone followed the action, both decision and action estimates shifted toward it. Spatial proximity also modulates early RP slope, with a steeper buildup in near versus far space, suggesting facilitated motor preparation within PPS. It further enhances the late RP amplitude when participants focused on their intention to act. Behavioural results show that actions are perceived as occurring earlier when the clock is displayed near compared to far, indicating that PPS influences the temporal perception of action timing. Overall, these findings highlight the dynamic interplay among external spatial context and internal cognitive processes in shaping motor preparation and action awareness. Importantly, a temporal internal attentional focus on intention to act modulates early RP—traditionally considered an unconscious stage of neural readiness. These results contribute to a deeper understanding of how PPS and the locus of attention on specific action phases affect action preparation and awareness, with potential implications for future research on the sense of agency and voluntary action decision making.

## Introduction

Voluntary actions are often referred to as self-initiated actions that are produced without any immediate external incentives, such as rewards or impending threats, and are instead initiated endogenously (Haggard, 2008, 2017; Passingham et al., 2010). This self-generation aspect is a critical facet of conscious control and the initiation of our actions. Notably, it has been consistently shown that these self-initiated actions are preceded by the Bereitschafts-potential (BP), also known as the “readiness potential” (RP), which is a slow build-up of negative EEG potential originating from pre-motor cortical areas (Kornhuber & Deecke, 1965), starting up to two seconds prior to self-initiated movements (Kornhuber & Deecke, 1965; Shibasaki & Hallett, 2006). The RP was first reported by Kornhuber & Deecke (1965) and later gained great attention through Libet et al.’s famous study, which claimed that the onset of the RP occurred several hundred milliseconds prior to the reported timing of the awareness of the decision to move (Libet et al., 1993; Libet et al., 1983). The timing of this decision was obtained by asking participants to report the time on a rotating clock when they first became aware of their intention to move (referred to as the will-time or W-time). Libet’s study suggested that the RP precedes conscious awareness of movement intention, igniting debates about its role in volitional control (Schmidt et al., 2016; Schurger et al., 2021; Triggiani et al., 2023).

The RP is commonly characterized as having an early and a late component based on the scalp distribution and the gradient of their negative potentials (Oken & Phillips, 2009; Shibasaki & Hallett, 2006). The early component (early RP, ∼2000 ms to ∼500 ms prior to movement onset) is slow and prolonged, with a gradual increase in negativity, which is localized symmetrically in the bilateral supplementary motor area (SMA) and premotor cortex, and was thought to represent the more general preparation for the forthcoming movement. In contrast, the late component (late RP) has a much steeper slope generated by activity in the contralateral primary motor cortex (M1), starting around ∼500 ms prior to movement onset (Shibasaki & Hallett, 2006).

Numerous studies have shown that the amplitude of the RP is influenced by various factors, which differ according to experimental paradigms. For instance, RP amplitude increases when participants consciously perceive themselves as preparing to move. This has been shown when participants made self-paced movements and reported their intention status in response to random visual probes (Parés-Pujolràs et al., 2019; Schultze-Kraft et al., 2020). However, recent findings employing alternative probing methodologies suggest that the RP might not reliably reflect the emergence of conscious intention, highlighting ongoing debates about its precise relationship to conscious volition (Gavenas et al., 2025; Parés-Pujolràs et al., 2023). Additionally, RP amplitude is greater in self-initiated actions compared to externally cued movements (Takashima et al., 2018) and in freely chosen actions compared to externally guided selections (Takashima et al., 2020).

Further, prior work has shown that when participants attempted to replicate the interval between two tones using two consecutive button presses, the early component of the RP (1600-800 ms before movement onset) was significantly larger preceding the second action which required attention to movement timing—unlike the first action, whose timing was incidental (Baker et al., 2012). Recent neurophysiological studies showed that the neural mechanisms underlying the RP are integrated with action timing processes, suggesting a critical role for RP in the temporal organization of voluntary actions (Emmons et al., 2017; Fifel, 2018). Moreover, predictive sensory outcomes of intended actions have been shown to amplify RP magnitude, supporting the view that RP reflects the integration of motor preparation and sensory prediction (Gärtner et al., 2025; Reznik et al., 2018; Vercillo et al., 2018; Wen et al., 2018). Beyond motor preparation, RP has been identified to encompass decision-related, non-motoric cognitive processes, further complicating interpretations of its functional role (Alexander et al., 2016; Schmidt et al., 2016). Thus, findings about how conscious intention, attentional allocation, and sensory outcome prediction influence the RP across studies employing different paradigms remain heterogeneous and sometimes conflicting (Gavenas et al., 2025; Triggiani et al., 2023). Moreover, previous studies have typically examined attentional focus using isolated manipulations of timing or outcome prediction, often in contexts where action reports were not required. In contrast, the present study introduces an integrative approach by examining how attentional focus (toward decision vs. action), and anticipated action outcomes interact to shape both RP dynamics and subjective action awareness. We hypothesize that directing attention towards the reporting of the decision or the execution of an action, will modulate the RP. Specifically, we anticipate that decision-focused attention will enhance the early phases of the RP, while action-focused attention will augment the later phases. Anticipated outcomes are also expected to influence the RP, reflecting the brain’s predictive mechanisms (Polezzi et al., 2008; Reznik et al., 2018).

Despite this extensive research on the modulation of the RP, one important dimension that has not been explored is the external spatial distance of the visual objects with which we interact. Spatial proximity, termed here as peripersonal space (PPS), i.e., the space immediately surrounding our body, is known to influence decision-making processes, urgency to act and the awareness of the voluntary action (Blini et al., 2018; Brozzoli et al., 2012; Brozzoli et al., 2011; Dureux et al., 2021; Kong et al., 2024; O’Connor et al., 2021; Patané et al., 2020). PPS constitutes a body-part-centered multisensory–motor reference frame that enhances cortical excitability within premotor and parietal regions, even in the absence of overt reaching or grasping movements (Brozzoli et al., 2012; Bufacchi & Iannetti, 2018). Despite its relevance to action execution, the role of PPS in voluntary action preparation and awareness remains underexplored. Rewarding objects located within the PPS has been shown to elicit more impulsive actions, even when no physical interaction is required, that is, when the physical action itself (keypress) is spatially invariant (O’Connor et al., 2021; O’Connor et al., 2014). A recent study further showed that objects’ proximity, namely positioning a Libet’s clock (Libet et al., 1983) with neutral valence within the PPS, led to earlier voluntary movement initiation (Kong et al., 2024), suggesting increased motor urgency, in line with converging evidence (O’Connor et al., 2021). This proximity also affected the subjective awareness of both the decision to act and, to a larger extent, the execution of the action (Kong et al., 2024). These behavioral findings highlight the need to investigate whether such spatial proximity effects extend to the neural level—specifically, whether they modulate the RP.

The present study addresses this gap by examining how spatial proximity, in addition to attentional focus and anticipated action outcomes, influences RP dynamics and action awareness. In this context, we employ a modified Libet clock within a virtual reality setup to evaluate the RP as participants interact with visual stimuli (the clock) located within or outside their PPS. Importantly, our paradigm allowed for the simultaneous manipulation of this spatial factor and the temporal attentional focus within the same voluntary action task—a design not previously explored in RP research. We hypothesize that spatial distance may also modulate the RP, with objects closer to the body potentially eliciting stronger preparatory responses. Several mechanisms could underlie such enhanced neural activity. First, stimuli within the PPS may increase response urgency, potentially but not necessarily translate into shorter response times, as PPS may modulate higher-order preparatory and predictive processes rather than overt motor execution. Alternatively, objects presented within PPS may influence the RP independently of urgency, by modulating the level of awareness during action preparation. Although the influence of awareness level on the RP is currently debated (Gavenas et al., 2025; Parés-Pujolràs et al., 2019; Parés-Pujolràs et al., 2023), PPS-related enhancement of RP amplitude could reflect increased access to—or integration of—awareness-related processes during motor preparation rather than changes in behavioral motor urgency per se. Finally, it is also possible that PPS benefits from a privileged neural representation for both perception and action, independently of urgency or awareness.

Overall, this investigation seeks to provide new insights into the features that shape the RP during voluntary actions, enhancing our understanding of the relationship between spatial factors, internal cognitive processes and neural activity.

## Methods

### Participants

Twenty-five healthy participants (14 females, age = 25.3 ± 3.9 years) were recruited for this study. We chose a target sample size of 25 participants based on prior EEG studies using similar within-subject designs to investigate readiness potentials with temporal binding or action-outcome manipulation (e.g., Jo et al., 2014; Reznik et al., 2018; Alexander et al., 2016), which typically include 20–30 participants. At the achieved n = 25, a sensitivity analysis shows we could detect effects of f = 0.24 (partial η² = .055) with α = .05 and power = .80. This target justification follows the practice guidance (Lakens, 2022). All participants were right-handed and unaware of the study’s aims and reported normal or corrected-to-normal vision, normal hearing, and no history of neurological disorders. Informed consent was obtained from each participant before their involvement in the study. They received a compensation of €30 for their participation. The study procedures were approved by the ethics committee (CEEI/IRB Comité d’Evaluation Ethique de l’Inserm, n°21-772, IRB00003888, IORG0003254, FWA00005831). The study adhered to the ethical standards of the Declaration of Helsinki.

### Material and apparatus

We adapted the Libet clock paradigm (Haggard et al., 2002; Libet et al., 1983) to a VR setup (Kong et al., 2017; Kong et al., 2024) to obtain stereoscopic vision of the clock. Participants sat comfortably and wore a head-mounted display (HMD, Oculus Rift CV1, resolution 2160 × 1200) while performing the task **(Fig. 1A)**. The virtual environment included a virtual room with two long, high, squared walls spaced 1.84 m apart, providing depth perception, and a virtual clock face **(Fig. 1B)** that participants were asked to fixate on. The clock face was marked with conventional “5-min” intervals, and had a ‘hand’ in the form of a green dot, which rotated clockwise with a period of 2560 ms per circle. The clock was presented randomly either at a distance close to (45 cm) or far from participants (3 m). The diameter of the near clock was 0.24 m, and the diameter of the far clock was 1.8 m, with both clocks having a visual angle of 29.86 degrees. Thus, the retinal size of the clock in the near and far space was corrected to avoid any potential size-dependent confound, as done in previous studies (Blini et al., 2018; Dureux et al., 2021; Kong et al., 2024; O’Connor et al., 2021). Before beginning formal testing, participants were asked to estimate the distance of the near and far clocks from themselves. The average estimations confirmed that perceived distances aligned with our intended presentation: near (Mean = 0.39 m, SD = 0.26 m) and far (Mean = 2.26 m, SD = 0.93 m).

**Figure 1.**
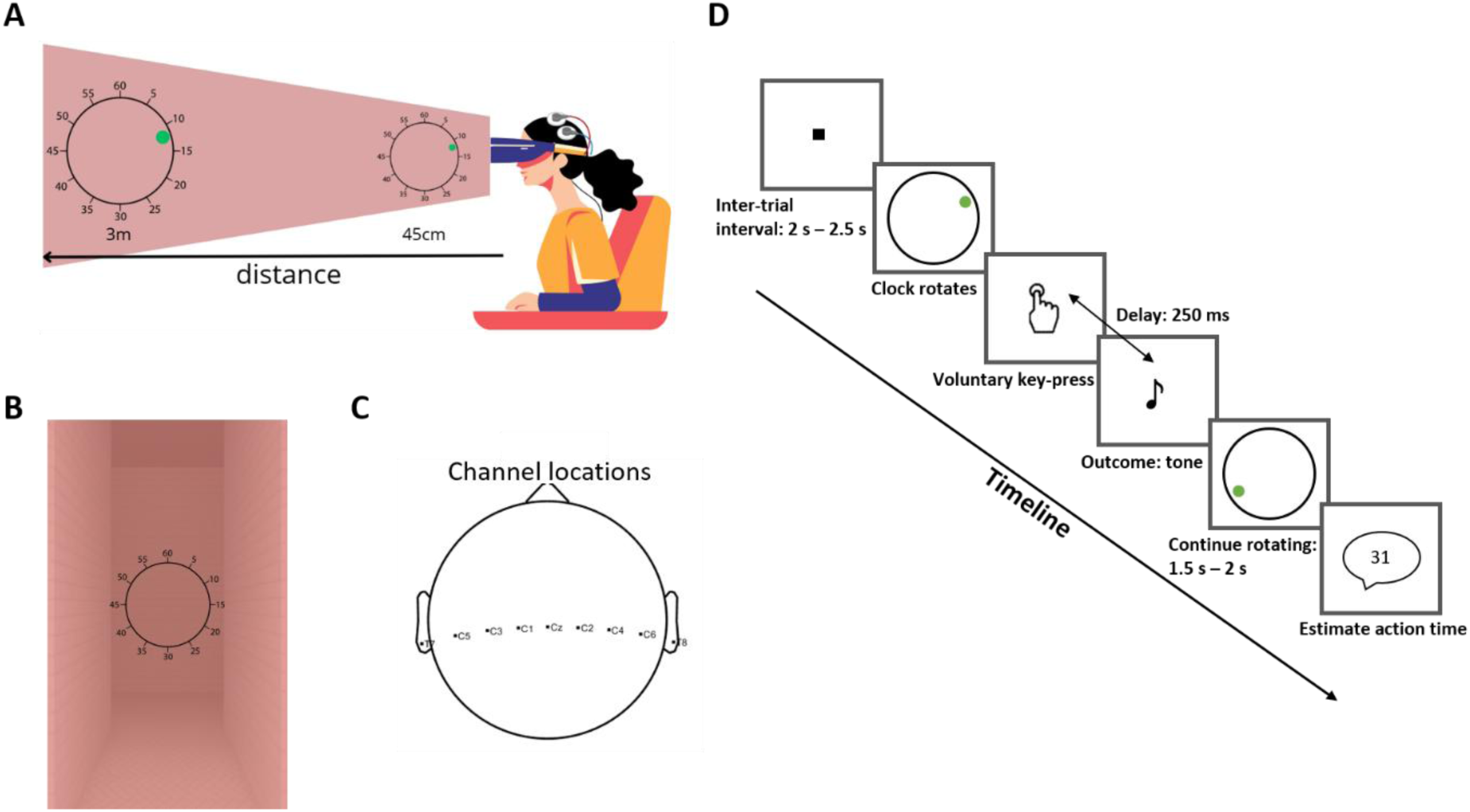
Experimental setup and schematic illustration of the task. (A) Illustration of the experimental setup: participants wore a VR headset for immersive display of the visual stimuli (rotating clocks) presented either in near space (45 cm) or far space (3 m). Clocks were rendered with size scaling to ensure identical retinal size across distances. Participants also wore an EEG cap with active electrodes, and surface EMG was recorded from the right extensor digitorum muscle to allow offline detection of movement onset. (B) Example of the virtual environment from the participant’s perspective, depicting a rotating clock embedded in a 3D-rendered corridor scene. (C) Top view of the EEG electrode configuration showing the nine central channels: T7, C5, C3, C1, Cz, C2, C4, C6, and T8. (D) Schematic representation of the trial structure in the Action Operant task. Each trial began with a fixation cross (2–2.5 s), followed by the appearance of a rotating clock. Participants made a voluntary keypress, which was followed by a 250 ms delay and an auditory tone (outcome). The clock continued rotating for another 1.5–2 s, after which participants verbally reported the position of the clock hand at the moment they believed they acted.

The key press was performed on a computer keyboard with the participants’ right index finger at a time of their own choice. The action consequence, when there was one, was an auditory tone (1000 Hz, sampling rate 44.1 kHz, 30ms duration, starting 250 ms after the keypress) played over the built-in headphone of the Oculus. During the inter-trial interval, a fixation square was displayed at the centre of the upcoming virtual clock and at the middle distance between the near and far space. The visualizations were programmed using the Unity platform (Unity 2018.4.22f1 Personal) and Microsoft Visual Studio 2019.

### EEG and EMG recording

EEG data were continuously recorded from nine scalp sites (T7, C5, C3, C1, Cz, C2, C4, C6, and T8) using active electrodes (ActiCap, Brain Products) attached to an EEG cap (EASYCAP GmbH), positioned according to the extended international 10/20 system **(Fig. 1C)**. The signal was recorded with a sampling frequency of 2500 Hz using Vision Recorder (Vision Recorder. Ink). The ground electrode was placed on Fpz, and all electrodes were online referenced to the right ear lobe. For stimulus presentation and behavioural response recording, Unity software was employed on the experimental control computer, which was synchronized with the EEG recording system. Event information for each trial was transmitted to the EEG system using a parallel port, ensuring accurate alignment of behavioural and EEG data. Electromyogram (EMG) signals were bipolarly recorded from the extensor digitorum muscle in the forearm via surface electrodes (Ambu® Neuroline 700), facilitating the precise temporal detection of the finger’s movement onset.

### Stimulus and experimental design

Our experiment consisted of a series of separate tasks unified by a commonly adapted Libet’s paradigm, in which a clock was pseudorandomly presented on each trial either near or far from the participant (Table 1). At the core of this paradigm was the observation of a rotating clock hand (**Fig. 1B**), with participants engaging in various tasks that involved key presses and/or auditory tone **(Fig. 1D)**. The primary difference between tasks lay in the specific event that participants were required (or not) to report. Each participant took part in all tasks, following a within-subject design.

**Table 1.**
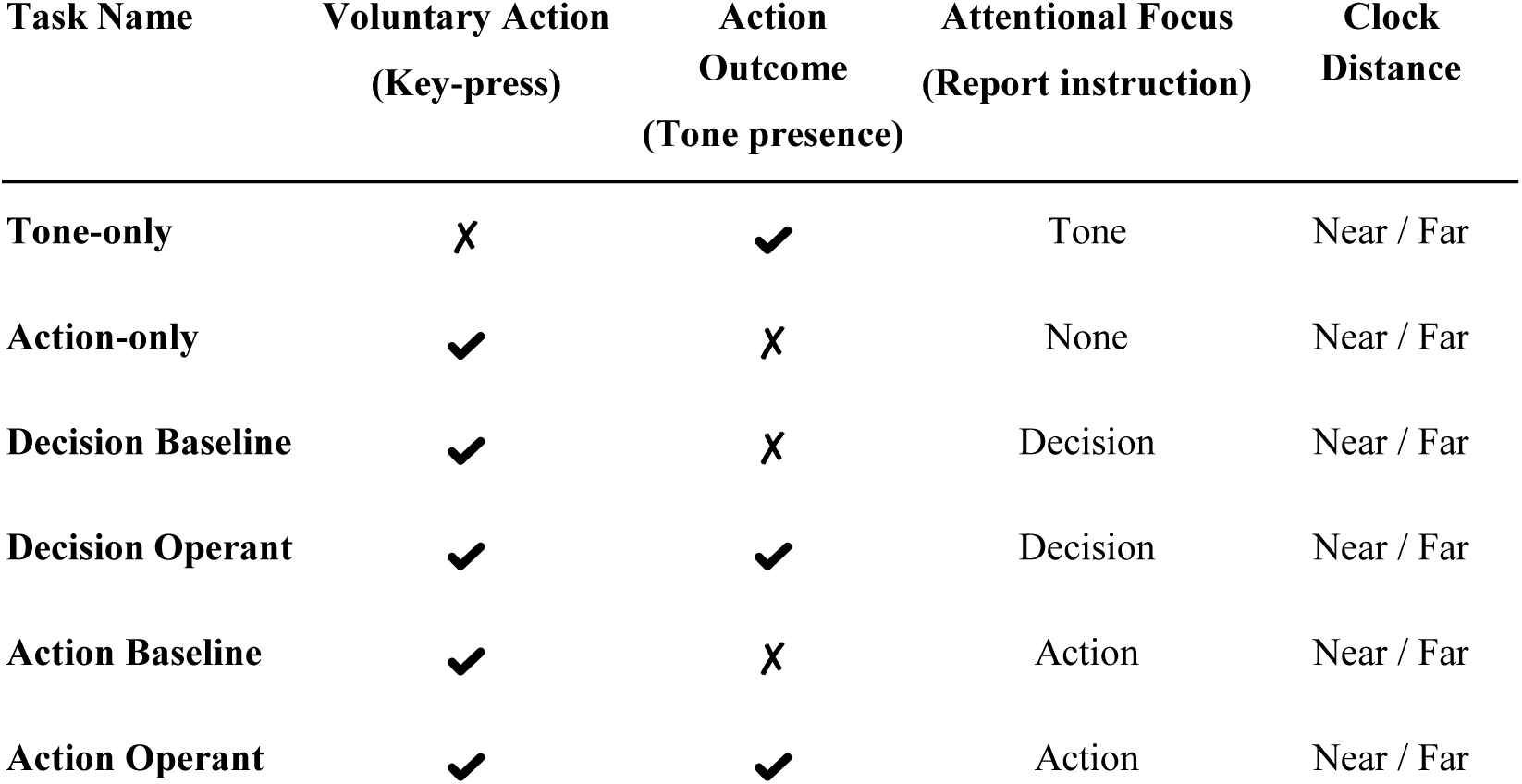
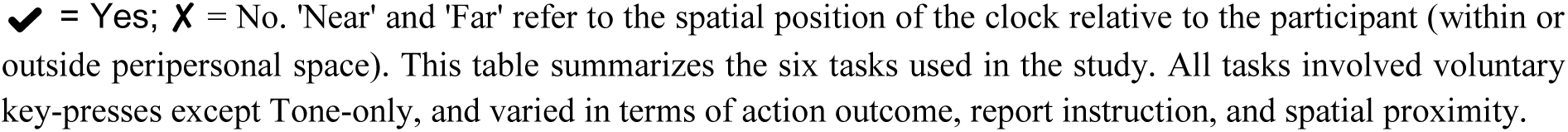
Overview of the experimental tasks.

| Task Name | Voluntary Action<br>(Key-press) | Action Outcome<br>(Tone presence) | Attentional Focus<br>(Report instruction) | Clock Distance |
| --- | --- | --- | --- | --- |
| Tone-only | ✗ | ✓ | Tone | Near / Far |
| Action-only | ✓ | ✗ | None | Near / Far |
| Decision Baseline | ✓ | ✗ | Decision | Near / Far |
| Decision Operant | ✓ | ✓ | Decision | Near / Far |
| Action Baseline | ✓ | ✗ | Action | Near / Far |
| Action Operant | ✓ | ✓ | Action | Near / Far |

✓ = Yes; ✗ = No. 'Near' and 'Far' refer to the spatial position of the clock relative to the participant (within or outside peripersonal space). This table summarizes the six tasks used in the study. All tasks involved voluntary key-presses except Tone-only, and varied in terms of action outcome, report instruction, and spatial proximity.

**Tone-only task:** Here, participants did not perform any key press. Instead, they were asked to report the timing of a tone that was randomly played by the computer, occurring 3 to 4.5 seconds after the onset of each trial (e.g., the onset of clock rotation). The experimenter then input their reports with another computer keyboard and did so for all other tasks that required participants’ reports (i.e., all tasks except the Action-only task). This single-event task provides a context to contrast against the other single-event task below (Action-only task) that only involves active movements.

**Action-only task:** After the clock hand started rotating, they could press the ‘zero’ key on a computer keyboard using their right hand index finger when they felt the urge to do so; in other words, at a time of their own choice. At odds with previous studies, to assess potential variations in the natural buildup of urgency, participants here were not instructed to wait until the clock rotation had completed at least one cycle. We reasoned that instructing participants to wait for at least one full cycle of clock rotation, thereby preventing them from acting as soon as they wished, could mask subtle differences in urgency between tasks. Indeed, our results show that many participants did not, on average, wait for a full cycle before pressing the key (**Supp. Fig. S2**). Once they pressed the key, the clock kept on rotating for a random duration (1.5 to 2s) before disappearing. This Action-only task sets the basis for simple action initiation. The instructions for pressing the key and the rotation of the clock were the same for all tasks with a key press (i.e., all tasks except Tone-only task, see above).

**Decision baseline task:** building upon the Action-only task, participants were additionally asked to verbally report at the end of the trial where the rotating hand was when they first became aware of their decision to press the button — referred to as “W-time” in Libet experiments (Libet et al., 1983). **Decision operant task**: similar to the Decision baseline, but here, a tone followed the key press after a 250 ms delay, allowing us to study the effect of predictable auditory consequences on decision reporting.

**Action baseline task:** participants estimated the timing of their key press, focusing on action timing. **Action operant task**: mirroring the Action baseline, but including a 250 ms delayed tone post-key press, to explore the influence of predictable auditory consequences on the reporting of action execution **(Fig. 1D)**.

These tasks collectively allowed us to investigate the influences of peripersonal space, temporal attentional focus, and action outcomes on the RP. For all tasks, a fixation square was presented during the inter-trial interval with a random duration between 2s and 2.5s, followed by the appearance of the virtual clock and the initiation of the clock’s rotation **(Fig. 1D)**. Compared to our recent study (Kong et al., 2024), this longer inter-trial interval was chosen in order to obtain a stable and noise-free baseline for EEG signals. For each task, 50 trials were performed for the near and far spaces, respectively, in a random order, in line with previous studies showing that the RP is best identified through averaging a large number (>30) of trials (Schurger et al., 2021). Notably, the distance at which the clock was displayed (45 cm or 3 m) was task-irrelevant **(Fig. 1A)**. Participants underwent 2 trials for familiarization before the formal testing in each task. The Tone-only was always the initial task, with all other tasks presented in a counterbalanced order.

### Data analysis

#### EMG analysis: Semi-automatic detection of movement onsets

For each trial, a high-pass Butterworth filter with a 10 Hz cutoff was applied to the EMG signals. EMG onset was then semi-automatically detected as follows (See also **Supp. Fig. S1** for the illustration of the EMG onset detection method). The signal was first transformed into its envelope. Then, the Peak was detected as the local maxima within the [350-0 ms] window preceding the computer-registered keypress. The baseline was calculated as the average signal over the period from 100 ms to 200 ms after the trial onset. The onset of the movement was defined as the time, within the [−350 0 ms] before the keypress, when the EMG envelope signal exceeded 50% of the Peak-Baseline difference. To augment the precision of our method, we integrated an interactive graphical user interface (GUI). This GUI enables manual marking of EMG activity onset, providing the flexibility for the onset to be identified even outside the predefined time window. This blend of automated processing and manual input allows for a more accurate determination of muscle activity onset.

#### EEG analysis: Readiness potential (RP)

EEG data were preprocessed and analysed using custom Python scripts and the MNE package version 1.5.1 for Python (Gramfort et al., 2013). Each task of each participant was preprocessed separately as follows: continuous EEG was filtered with a 0.1 Hz high-pass filter, a notch filter at 50 Hz, and a 30 Hz low-pass filter, followed by artifact correction using the *Autoreject* algorithm (Jas et al., 2017). This algorithm automatically detects and corrects transient artifacts on a trial-by-trial basis using data-driven thresholds and local interpolation. This approach minimizes the influence of ocular and other transient artifacts without requiring manual ICA-based correction. EEG data from the Cz channel were epoched between −1.2 and 0.1 s around either the computer-recorded keypress, or the EMG movement onset. For the Tone-only task, EEG data from the Cz channel were epoched between −1.2 and 0.1 s around the tone onset. Additionally, epochs were excluded from analysis if they showed a peak-to-peak (max–min) amplitude exceeding 100 μV within the −1.2 to +0.1 s window. This criterion was chosen, instead of rejecting epochs containing sample points exceeding fixed thresholds, in order to avoid unnecessary rejection of epochs due to slow drifts/DC offsets. Visual inspection of the EEG traces revealed brief transient deflections shortly before action in some tasks. We inspected the data to ensure that these were not eye-movement or blink artifacts. This effect was largely reduced when epochs were time-locked to EMG-detected movement onset. Our interpretation is that some participants lifted their finger before pressing the key, producing a minor pre-movement artifact. Importantly, this did not affect the overall RP pattern or the main conclusions of the study.

Given that we do not know when volition begins, we cannot determine the appropriate baseline for analyzing the RP (Haggard, 2024). Any pre-movement interval may already include neural buildup related to voluntary preparation. Subtracting such activity could artificially distort the slow, gradual dynamics characteristic of the RP. Additionally, in light of the concerns that baseline correction may artificially influence RP amplitude differences—potentially leading to misleading results (Delorme, 2023; van Driel et al., 2021), we opted to analyze and report our data without applying baseline correction. Although we intentionally did not apply a traditional baseline correction however, the use of a high-pass filter effectively centers the continuous data around zero over time. Thus, with a sufficient number of trials, the mean voltage should approach zero, functionally mimicking a baseline correction without imposing a specific pre-movement reference window. This rationale is empirically validated by our Tone-only condition (see results), in which no voluntary movement occurred: the mean signal remained around zero, confirming that the pre-processing procedure provided a stable reference and did not introduce systematic drifts.

The early RP was quantified as the mean amplitude during the first 600 ms [−1200 to −600 ms], while the late RP was the mean amplitude of the following 600 ms [−600 to 0 ms]. In addition to analysing the mean amplitudes, we also examined the slopes of the early and late RP components, as slopes are less affected by baseline correction issues. The early RP slope was computed as the difference between the average voltage in the first 50 ms of the early RP window (−1200 to −1150 ms) and the average voltage in the last 50 ms of the same window (−650 to −600 ms), divided by the duration of the early RP interval. The late RP slope was computed analogously, by subtracting the average of 50 ms preceding the late RP component [−650 to −600 ms] from the average of the last 50 ms [−50 to 0 ms]. This difference was then normalized by the 600 ms duration, representing the rate of change in the late RP signal (Jo et al., 2014).

A three-way repeated measure ANOVA with balanced factors 2 (distances: near vs. far) × 2 (report instructions: decision vs. action) × 2 (tone presence: baseline vs. operant) was conducted separately for the early RP, late RP, early RP slope and late RP slope. Additionally, a two-way repeated measures ANOVA with factors 2 (Distance: near vs. far) × 2 (Task: action vs. tone) was performed for Tone-only and Action-only tasks. One sample t-tests were also conducted for the Tone-only and Action-only tasks, respectively, to further assess whether the typical RP profile appears only in the action task rather than in response to an external stimulus.

#### Subjective awareness analysis

To investigate the impact of distance on the subjective awareness of action decision, execution, and the existence of temporal binding induced by the presence of a tone, we considered all tasks with a report (i.e., all tasks except the Action-only task) and we calculated time estimates as follows. First, we converted the reported position of the clock in milliseconds. Then, for the action report and tone report tasks, the actual time of the keypress (or the tone, respectively) was subtracted from the subjective reported time of the action (or the tone, respectively). These estimated times thus correspond to action and tone judgment errors. The values were negative when the event (action or sound) was subjectively perceived before the actual event. Similarly, for the decision report tasks, the time of the keypress was subtracted from the subjective reported time of the decision. The values were negative when the decision was subjectively perceived before the actual action. A 2 × 2 × 2 repeated measure ANOVA was performed on the estimated time with the factors clock distance (near, far), report instruction (decision, action), and tone presence (baseline, operant). Additionally, a paired t-test was carried out for the Tone-only task.

#### Movement initiation time (MIT) analysis

To explore how proximity influences action initiation time, we calculated the mean movement initiation time across trials for each task that involved a keypress (i.e., all tasks except the Tone-only task). This calculation entailed determining the time elapsed between the onset of the clock rotation and the computer-registered keypress (keypress movement initiation time or keypress MIT), or the initiation of movement as detected in the EMG recordings (EMG MIT). A paired t-test was conducted for the Action-only task to provide insights into the effects of the distance of the clock from the body on action initiation when participants had nothing to report. In addition, a three-way rmANOVA 2 (distances: near vs. far) × 2 (report instructions: decision vs. action) × 2 (tone presence: baseline vs. operant) was run on all other tasks. Both the paired t-test and the ANOVA were run separately on keypress MIT and EMG MIT.

None of the factors significantly influenced either the computer-registered keypress time or the EMG-detected movement onset time (see the mean results in **Supp. Fig. S2** and the distribution in **Supp. Fig. S3 and S4).** Moreover, the factors did not influence the motor times, which are the differences of MITs between keypress and EMG, either (all p > 0.21). However, we also conducted an unplanned exploratory analysis, in line with previous work showing that urgency can influence motor time in the context of speed–accuracy trade-offs (Steinemann et al., 2018). In our data, the distribution of motor times revealed that most values fell within the critical range of 0–0.1 s, but interestingly, a second cluster appeared between 0.1 and 0.5 s (note that motor times rarely exceeded 0.5 s, only in 0.1%–1.4% of trials on average; see **Supp. Fig. S5**). To further characterize the dynamics of movement, the same three-way repeated measures ANOVA was thus applied on the percentage of motor times falling inside the 0.1-0.5s range. Note that the data of these percentages in the Action baseline and Action operant tasks were found to be non-normally distributed. We nevertheless proceeded with the ANOVA analysis as our experiment was explicitly designed as a factorial experiment and designed to test whether there were interactions: balanced ANOVA is quite robust to departures from normality (Montgomery, 2017). Post-hoc two-by-two comparisons were tested with both Wilcoxon signed-rank tests and t-tests for tasks in which there were normality violations. As the pattern of statistical significance was unchanged with both types of tests, only the latter is reported in this manuscript.

All the data analyses were carried out using customized MATLAB programs (2019b, Mathworks, Natick, MA) and Python 3.0. All statistical analyses were performed using the IBM SPSS statistical package (SPSS 29.0). The significance level was set at α = 0.05, and Bonferroni correction was applied for multiple comparisons.

## Results

### EEG results

To characterize the influence of clock distance, report instruction and tone presence on the shape of the RP in both epochs aligned on computer-recorded keypress **(Fig. 2)** and epochs aligned on EMG movement onset **(Supp. Fig. S6)**, we calculated the mean amplitudes for the early and late RP periods, as well as the slopes of early and late RP. The analyses on both types of epochs yielded consistent results. Therefore, for conciseness, only the detailed findings on the epochs aligned on computer-recorded keypress are reported **(Fig. 3)**. It should be noted that ‘larger’ metrics indicate greater negative values.

**Figure 2.**
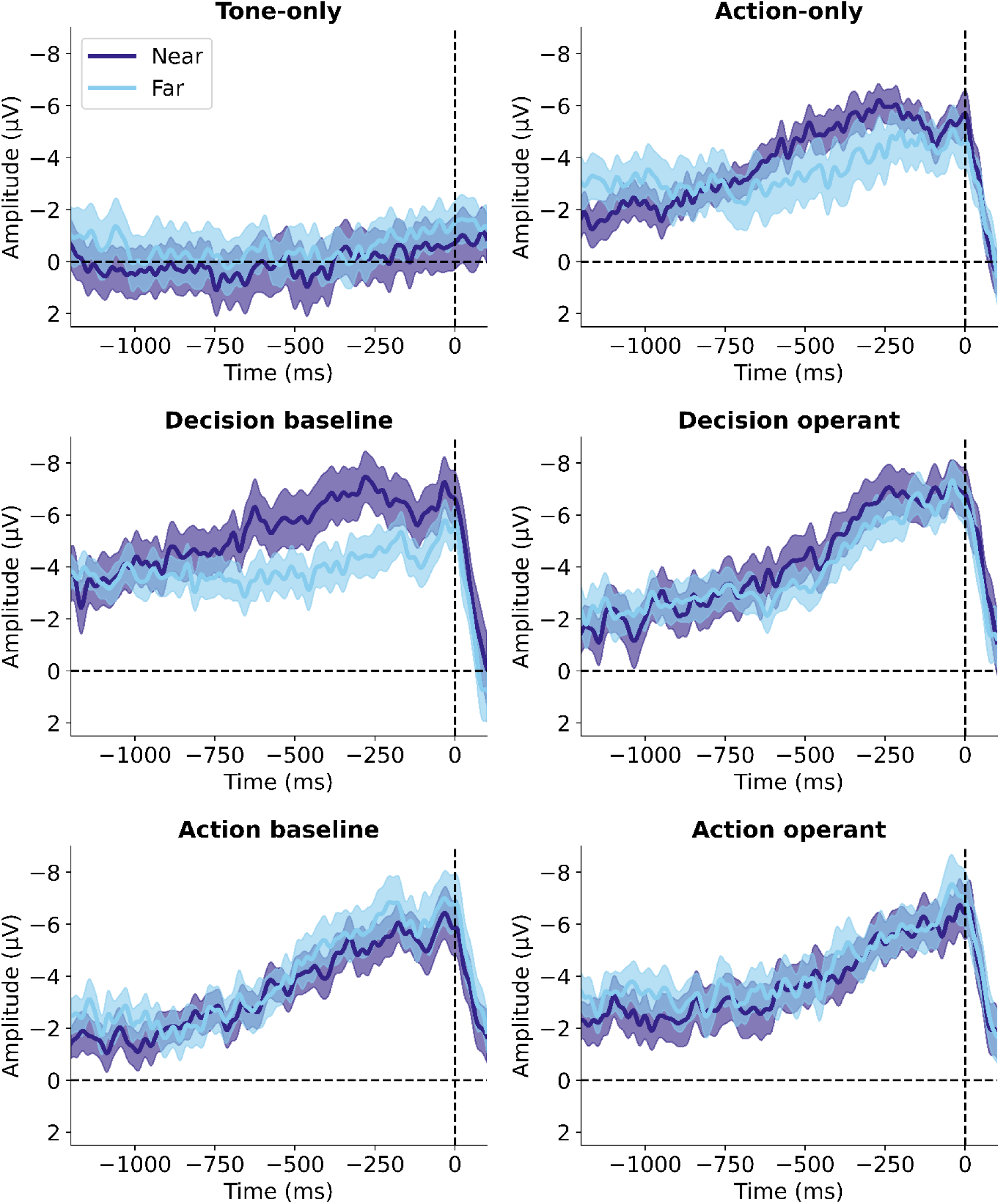
The time-course of readiness potential (RP) without baseline correction across different experimental tasks. (described in Table. 1). The vertical dashed line at time zero represents the computer-recorded keypress for the first five tasks or the tone onset for the “Tone-only” task. Light blue: clock presented near; Purple: clock presented far. The shaded regions surrounding each line represent one standard error of the mean.

**Figure 3.**
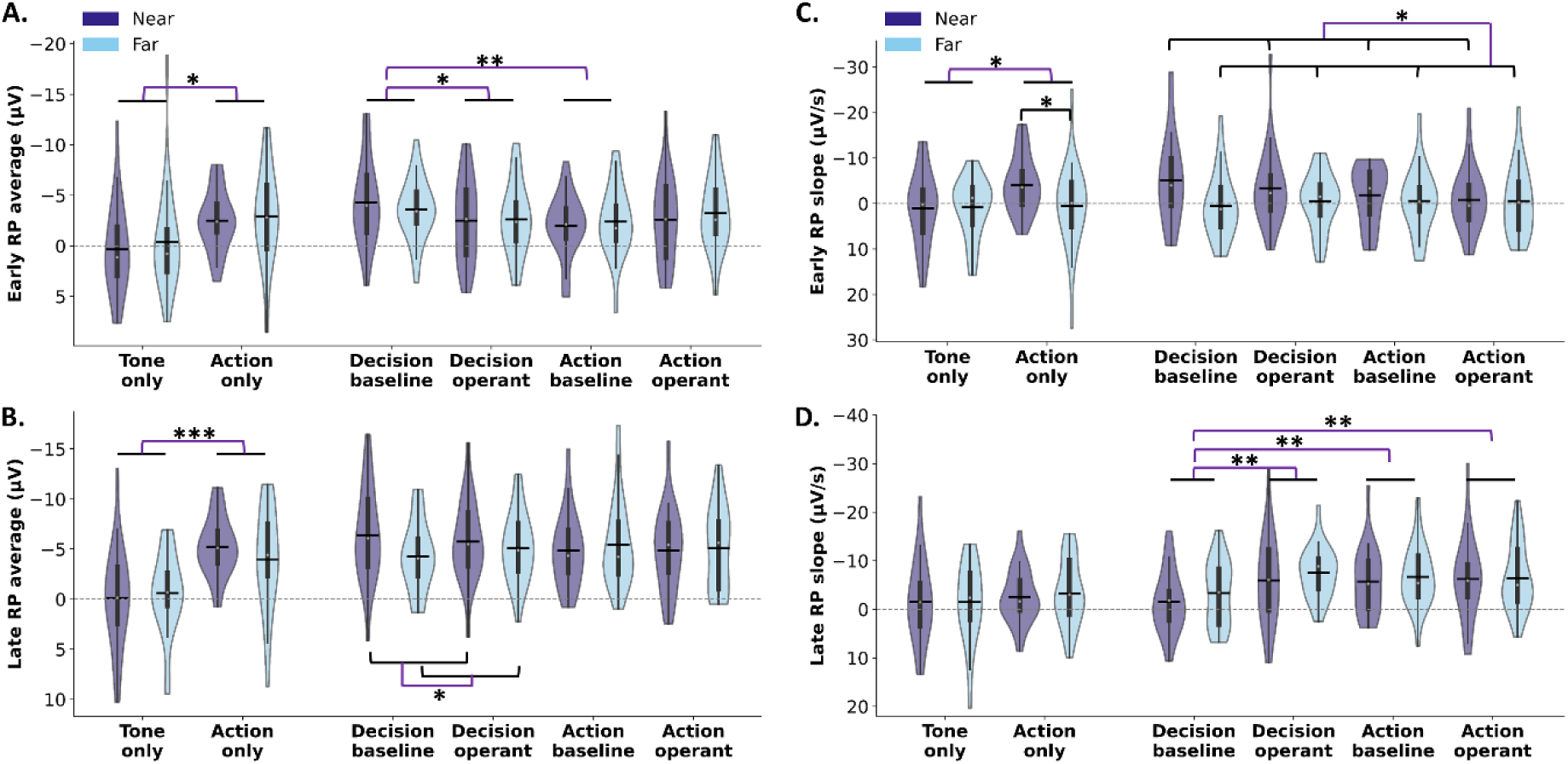
RP metrics for each experimental task (without baseline correction). **A.** Early RP mean amplitudes; **B.** Late RP mean amplitudes; **C.** Early RP slopes; **D.** Late RP slopes. Note that greater negative potentials represent a greater amplitude from 0 and that negative slopes indicate a steeper negative descent in potential. The ‘Near’ condition is represented by light blue violins, while the ‘Far’ condition is depicted by purple violins. The width of each violin represents the probability density of data points at different RP metrics. The white dot within each violin represents the median RP metrics, the thick vertical line represents the interquartile range, and the horizontal black bar represents the mean RP metrics within each task. ** for p < 0.01, * for p < 0.5.

The two-way repeated measures ANOVAs were conducted on the mean amplitudes of the early and late RP, and on the early and late RP slopes for the Tone-only and Action-only tasks. A significant main effect of Task was observed for the early RP (F (1,24) = 4.68, p = .04, partial η² = .16), the late RP (F (1,24) = 19.90, p < .001, partial η² = .45), and the early RP slope (F (1,24) = 4.54, p < .05, partial η² = .16), with greater early and late RP amplitudes in the Action-only compared to the Tone-only task. No other main effects or interactions were found (p > .16). To further investigate the presence of the RP, one-sample t-tests were conducted against zero. In the Tone-only task, none of the RP components (early and late RP mean amplitudes, early and late RP slopes) differed significantly from zero (all p > .37), indicating the absence of a typical RP profile in the absence of movement. In contrast, almost all RP measures in the Action-only task were significantly different from zero (early RP mean amplitude: near p < .001, far p = .004; late RP mean amplitude: near and far p < .001; late RP slope: near p = .026, far p = .022), reflecting a robust RP signal. For the early RP slope, it was significantly greater than zero when the clock was presented in near space (p = .003), but not in far space (p = .77). And, the paired t-test between the two distances was significant (p = .026). These findings confirm that a reliable RP profile emerges only in the presence of voluntary movement, and is absent when no action is performed.

The three-way repeated measure ANOVA was conducted for the balanced tasks involving keypress and reports, with the factors clock distance (near, far), report instruction (decision, action), and tone presence (baseline, operant). For the early RP mean amplitude **(Fig. 3A)**, all the main effects were not significant (all p > .16), but the interaction between the report instruction and tone presence was significant (F (1,24) = 8.61, p = .007, partial η² = .26). Pairwise comparisons revealed that the early RP mean amplitude was significantly larger in the decision baseline task than in the decision operant task (Mean difference ± SE = −1.38 ± 0.67, p < .05, 95% CI [−2.76, −0.004]), and than in the action baseline task (Mean difference ± SE = −1.74 ± 0.49, p = 0.0015, 95% CI [−2.74, −0.74]). None of the other pairwise comparisons yielded significant results (all p > .08). No other significant interactions were found (all p > .25). These results indicate that the early RP amplitude was enhanced in the decision baseline task, possibly suggesting stronger neural engagement in early preparatory activity when participants attend to their decision without the expectation of an external outcome.

For the late RP mean amplitude **(Fig. 3B)**, all the main effects were not significant (all p > .23), but the interaction between the report instruction and clock distance was significant (F (1,24) = 7.70, p = .01, partial η² = .24). Pairwise comparisons revealed that in the decision reporting tasks, the late RP mean amplitude was significantly larger when the clock appeared in near space compared to far space (Mean difference ± SE = −1.36 ± 0.55, p = .02, 95% CI [−2.49, −0.23]). None of the other pairwise comparisons yielded significant results (all p > .15). No other significant interaction was found (p > .39). These results indicate that the modulation of the late RP amplitude by spatial proximity is specific to decision-reported tasks.

For the early RP slope **(Fig. 3C)**, the main effect of Distance was significant F (1,24) = 5.62, p = .026, partial η² = .19), suggesting the early RP slope was steeper when the clock was presented in the near space than far space. No other significant main effects or interactions were observed (p > .10).

For the late RP slope **(Fig. 3D)**, there were a significant main effects of tone presence (F (1,24) = 12.07, p = .002, partial η² = .34), and a significant interaction between report instruction and tone presence (F (1,24) = 6.25, p = .02, partial η² = .21). Pairwise comparisons revealed that the late RP slope was significantly shallower in the decision baseline condition than in the decision operant condition (4.29 ± 1.19, p = .001, 95% CI [1.84, 6.74]), than in the action baseline condition (3.78 ± 1.20, p = .004, 95% CI [1.32, 6.25]), and than in the action operant condition (3.88 ± 1.20, p = 0.004, 95% CI [1.40, 6.35]). None of the other pairwise comparisons yielded significant results (all p > .77). No other significant main effects or interactions were found (p > .13). These results suggest that the late RP slope becomes steeper in tone-present conditions, indicating a more rapid buildup of neural activity prior to movement execution. Additionally, the shallowest late RP slope was found in the decision baseline task, which mirrors the results reported for the early RP mean amplitude.

Note that no significant correlation was found between the mean movement initiation time and three of the RP metrics, including the early RP mean amplitude (adjusted p > 0.24), late RP mean amplitude (adjusted p > 0.4), and late RP slope (adjusted p > 0.1). For the correlation between mean movement initiation time and the early RP slope, a single significant correlation emerged in the decision baseline task when the clock was presented in the far space (adjusted p = .02). Given that this effect did not replicate across other tasks or spatial conditions, and considering the number of correlations tested, this isolated finding is likely to reflect statistical variability rather than a systematic relationship. Thus, we did not find evidence that movement initiation time related factors contributed meaningfully to the RP effects observed in our study.

### Subjective awareness of action decision and action execution

Except in the Action-only task, participants were required to report the estimated time of their decision, their action or just a tone played by the computer **(Fig. 4)**. The paired t-test for the Tone-only task showed no significant difference between the tasks in which the clock was near (Mean ± SE = 134.75 ± 8.84) and far (Mean ± SE = 133.50 ± 7.88 s, t (24) = 0.38, p = .71). This suggests that the distance of the clock from the body did not affect time perception itself when there was no voluntary movement involved.

**Figure 4.**
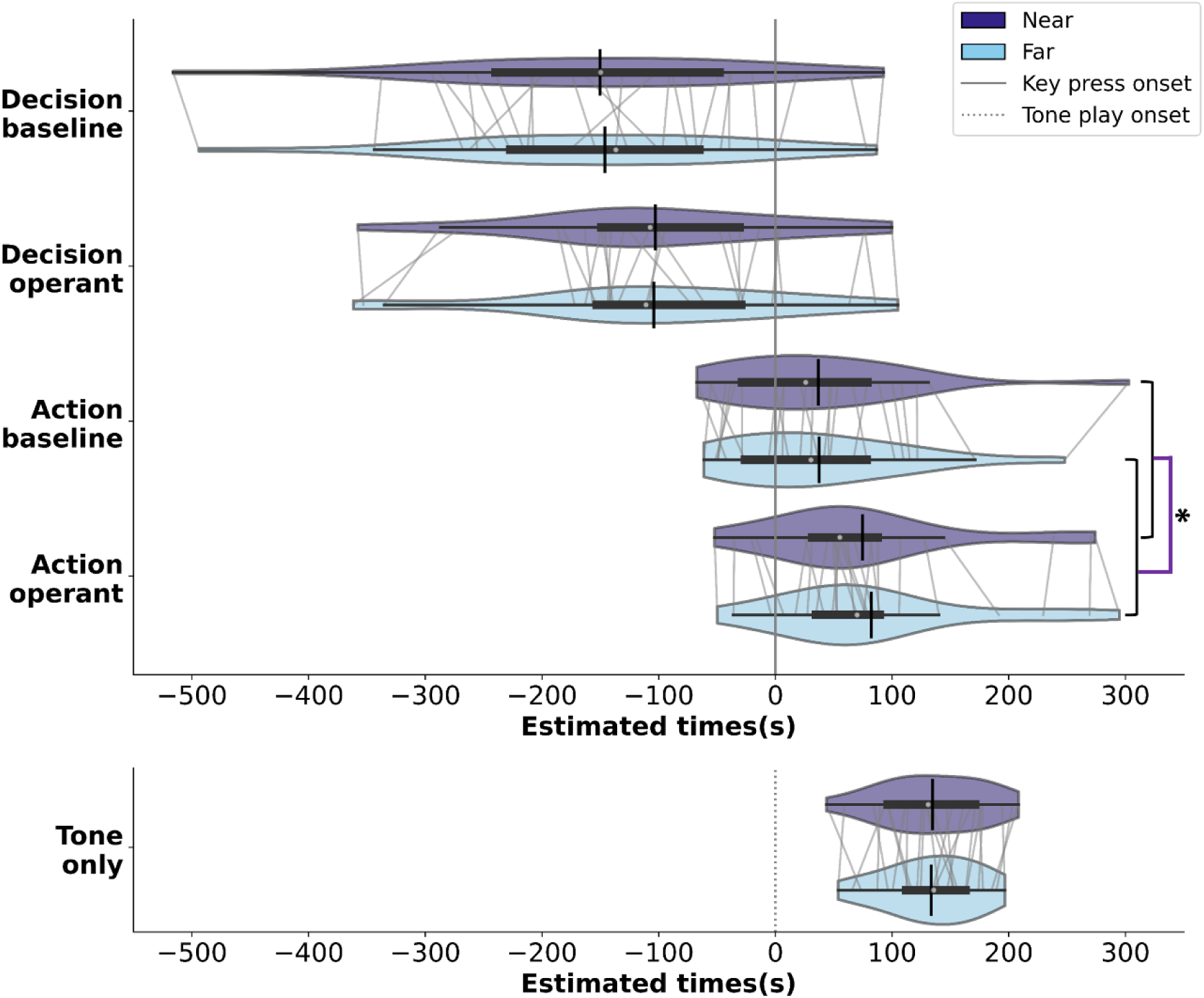
The distribution of estimated times for Decision estimates, Action estimates and Tone-only estimates. The light blue and purple violins distinguish between near and far distances, respectively. For the subplots of “Decision estimates” and “Action estimates”, the x-axis is centred around the key-press onset (a solid vertical line), showcasing estimated time variations in relation to key-press time. For “Tone-only estimates”, the x-axis is centred around the tone play onset (a dotted vertical line). The width of each violin represents the probability density of data points at different times. The white dot within each violin represents the median estimated times, the thick horizontal line represents the interquartile range, and the vertical black bar represents the mean estimated times within each task.

The 2 × 2 × 2 repeated measure ANOVA on the estimated time of tasks involving a keypress and a report, with the factors of distance (near, far), report instruction (decision, action), and tone presence (baseline, operant), did not show a significant main effect of distance (F (1,24) = 3.18, p = .087, partial η2= .12). The main effect of report instruction was significant (F (1,24) = 96.51, p < .001, partial η2= .80): as expected, participants perceived their decision as occurring earlier than their action. The interaction between distance and report instruction was not significant (F (1,24) = 0.66, p = .42, partial η2= .03). Nonetheless, aligning with our *a priori* hypothesis, based on our recent study (Kong et al., 2024), which suggested that distance effects may emerge when action and decision estimates are analysed separately, we conducted separate paired t-tests on action and decision estimates (after averaging across tone-presence conditions). Distance had a significant effect on action estimates (t (24) = −2.48, p = .02), but not on decision estimates (t (24) = −0.32, p = .75), suggesting that the action time was reported significantly earlier in the near relative to the far space.

Additionally, the ANOVA showed that the main effect of tone presence was highly significant (F (1,24) = 20.48, p < .001, partial η2= .46). This indicated that both the perceived times of decision and action were shifted towards the tone compared to when there was no tone. Thus, there was a temporal binding between the perceived time of action and its consequence, as well as between the perceived time of decision and the subsequent consequence. No other interactions were significant (p > .26).

### Movement initiation times and motor times

As mentioned in the data analysis section, neither the computer-registered keypress time nor the EMG-detected movement onset time was significantly influenced by any of the factors **(Supp. Fig. S2)**. However, the distribution of the difference between these two timings, which characterizes the motor time (from the beginning of muscle activity to the keypress) revealed an intriguing pattern **(Fig. 5)**. The three-way repeated measure ANOVA for the tasks involving a keypress and a report revealed significant main effects of distance (F (1,24) = 7.57, p = .01, partial η2 = .24), report instruction (F (1,24) = 12.07, p = <.01, partial η2 = .34), and tone presence (F (1,24) = 9.29, p = <.01, partial η2 = .28) on the percentage of trials with longer motor time (between 0.1 and 0.5 s). In addition, the interaction between distance and tone presence was significant (F (1,24) = 14.93, p < .01, partial η2 = .38). The interaction between distance, report instruction, and tone presence was significant (F (1,24) = 33.93, p < .001, partial η2 = .59), with significantly more trials for which the EMG activity started early (>0.1 s) before the keypress (i.e., longer motor time) when the clock was closer to the body compared to when it was far in the decision baseline task only (Mean difference ± SE = 11.58 ± 2.36, p < .001, 95% CI [6.71, 16.46]). None of the other pairwise comparisons yielded significant results (all p > .06). The other interactions were not significant (all p > .5). Finally, the paired t-test for the Action-only task did not show significantly more trials with longer motor time (> 0.1 s) when the clock was near the body compared to when it was far (t (24) = 1.45, p = .16).

**Figure 5.**
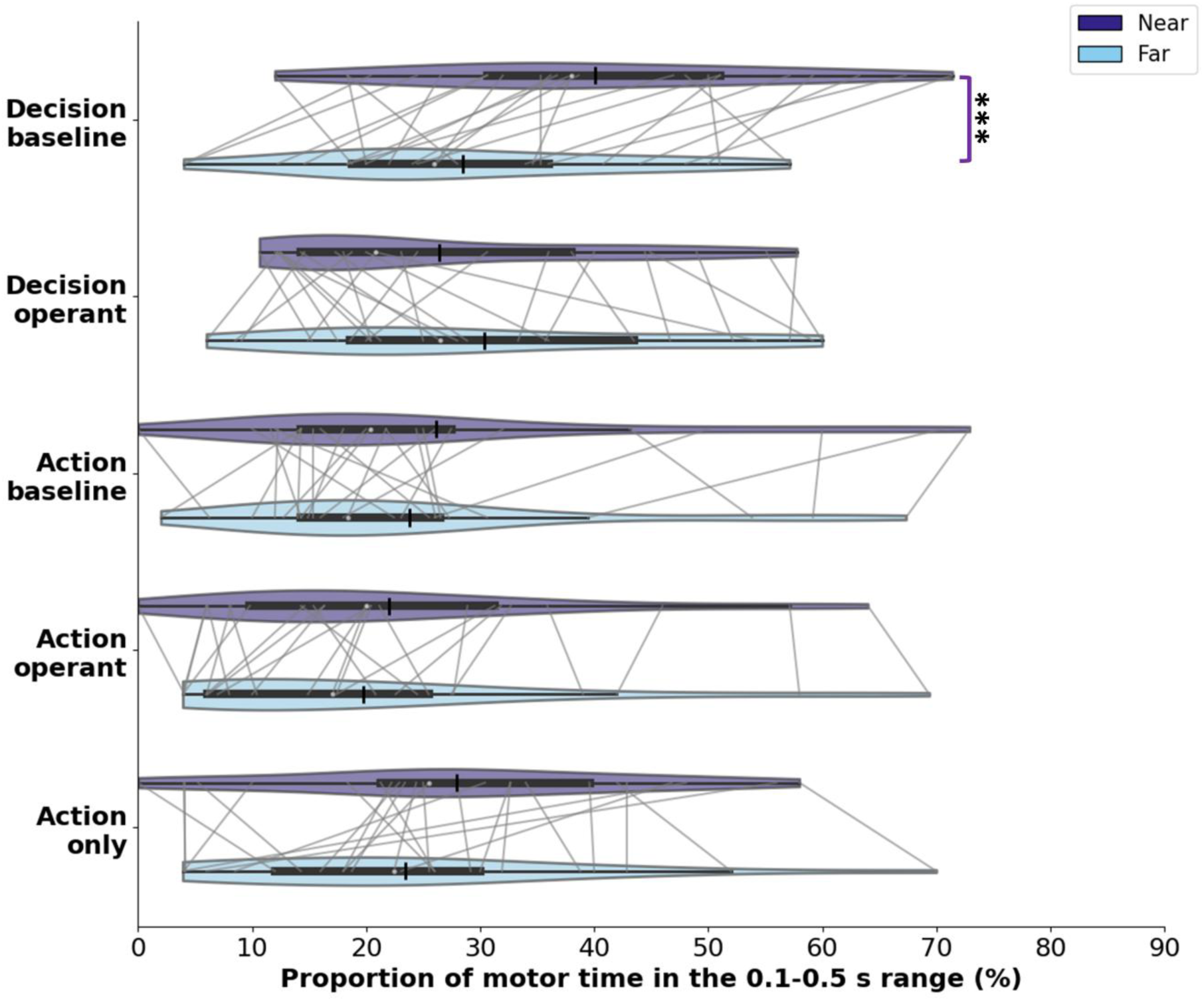
Proportion of motor times within the critical range of 0.1 to 0.5. **s** (note that motor times rarely exceeded 0.5 s). The light blue and purple violins distinguish between near and far distances, respectively. Each violin’s width reflects the probability density of data points across the percentage of motor times on the x-axis, offering a visual portrayal of data distribution within each task. The white dot within each violin represents the median movement time, the thick horizontal line represents the interquartile range, and the vertical black bar represents the mean motor time within each task. *** for p < 0.001.

## Discussion

This study investigates how spatial proximity, attentional focus, and anticipated action outcomes influence motor preparation and action awareness by exploring the impact of object distance, report instructions, and action outcomes on the readiness potential (RP), while also analyzing subjective temporal perception and movement time during voluntary actions. EEG and EMG data were collected as the participants interacted with a Libet clock that was presented either close to or far from their body, in a virtual reality setting. Our results showed that shifting the focus of attention to the earlier phase - when participants reported the timing of their intention to act - enhanced the early RP, suggesting a stronger neural engagement in early preparatory activity. Additionally, shifting the attention to later phases, either by focusing on the action or by the mere-relevance of a task-irrelevant tone presented shortly after the keypress also influenced the RP by increasing the late RP slope. In cases where the tone was present, this effect was accompanied by a shift of the subjective reports of decision and action to later timing - a well-known temporal binding phenomenon (Haggard et al., 2002). Spatial proximity also influenced RP dynamics. Specifically, the early RP slope showed a steeper buildup in near compared to far space. Moreover, spatial proximity also enhanced the late RP amplitude when participants focused on their intention to act. Behavioural findings further showed that when the clock was near, actions were perceived as occurring earlier compared to when it was far, suggesting a spatial influence on the temporal perception of action. EMG results showed that spatial proximity increased the likelihood of earlier-initiated and longer-lasting movements, particularly in decision-reporting tasks that lack action consequences. Below, we will discuss each of these main results in detail.

Focusing on decision timing (without subsequent action outcomes) rather than on action timing yielded a higher early RP amplitude and a less steep late RP slope, suggesting increased preparatory neural engagement during intention formation. In contrast, focusing on action execution leads to a steeper late RP slope, indicating a more rapid motor buildup when attention is directed toward movement onset. In the present study, the “early RP” window was defined as the interval from 1200 to 600 ms before the keypress. Since motor time was below 400 ms in most trials **(Supp. Fig. S3)**, this window typically ended more than 200 ms before movement onset, as determined with EMG. This places the early RP window within the typical window between the RP onset and the W-time from Libet et al. 1983, which is traditionally considered as an unconscious stage of movement preparation (Armstrong et al., 2022; Libet et al., 1983; Shibasaki & Hallett, 2006). However, our findings importantly suggest that this unconscious stage of neural preparation phases (i.e. the early RP) is significantly influenced by the attentional focus on the intention to act. This indicates that modifications in RP are influenced not just by motoric demands, but also by the nature of cognitive engagement required by the task, challenging the view that RP is predominantly an unconscious precursor to voluntary movement. Early RP is believed to reflect general preparation for the impending movement, involving bilateral activation of the Supplementary Motor Area (SMA) (Oken & Phillips, 2009), which plays an important role during motor preparation and in controlling perceptual processing during voluntary actions (Nachev et al., 2008). Additionally, SMA activity increases when individuals focus attention on their intent to move, rather than on the movement itself (Lau et al., 2004). This suggests that SMA and pre-SMA activity may represent the intention (Haggard, 2005), a hypothesis supported by our observation of increased early RP when participants focused on their intention timing. Future research should thus aim to consolidate and deepen these findings, particularly exploring whether there is enhanced pre-SMA activity in scenarios like our decision estimation task, without subsequent action outcomes.

In this study, the locus of attention was manipulated by the instructions (reporting decision time vs. action time), but also by the presence or absence of a tone (action-outcome) following the keypress, even though the tone was task-irrelevant. In this context, we replicated the temporal binding effect: when a tone was present, both decision (which has received much less attention in previous studies) and action estimates shifted towards the tone, as observed in our earlier work (Kong et al., 2024). Traditionally, temporal binding has been interpreted as a proxy of the sense of agency in voluntary movement (Haggard & Chambon, 2012). However, it also has been shown to result from somatosensory integration and temporal prediction (Kirsch et al., 2019; Gaiqing Kong et al., 2024; Suzuki et al., 2019), causal inference (Buehner, 2015; Buehner & Humphreys, 2009), and spatial attention drawn to the clock position at the time of the tone onset (Cao, 2024). In the present study, these decision-binding and action-binding effects were accompanied by an increase in the late RP slope, suggesting that anticipating an action outcome enhances the rate of neural buildup during the final stages of motor preparation. Interestingly, previous studies have also reported that predicted sensory outcomes can modulate either the amplitude or the slope of the RP (Gärtner et al., 2025; Reznik et al., 2018; Vercillo et al., 2018; Wen et al., 2018). The specific RP component affected may depend on factors such as the reliability and specificity of the predicted stimulus, the reporting instructions, and analytical choices in EEG preprocessing, including whether and how baseline correction is applied. Indeed, since baseline correction can influence RP amplitude measurements, it is important to recognize its potential impact on reported effects. As a validity check for our choice to omit baseline correction, we confirmed that the ‘Tone-only’ task, during which participants estimated the timing of an externally triggered tone, produced no typical RP profile, as expected. Overall, our results confirmed that the brain’s preparation for action is not only about motor execution, but also encompasses the anticipation of sensory outcomes, as evidenced by an increase in the late RP slope, reflecting an integrated sensory-motor process. Understanding this relationship could have significant implications for designing cognitive and motor rehabilitation programs, particularly in disorders where sensory feedback processing is impaired. Future research could explore how varying the type and timing of sensory feedback influences RP, potentially leading to more effective rehabilitation techniques that leverage sensory-motor integration.

Our findings further indicate that spatial proximity influences motor preparation dynamics. Specifically, the early RP slope was steeper in near compared to far space, suggesting facilitated motor preparation within PPS. In addition, spatial proximity enhanced the late RP amplitude when participants focused on their intention to act, indicating increased cortical excitability preceding voluntary movement. This enhanced motor preparatory activity is consistent with the notion that stimuli within PPS are prioritized by the brain due to their heightened relevance for bodily interaction and potential threat, thereby engaging sensorimotor systems more robustly (Blini et al., 2021; Bufacchi & Iannetti, 2018; Noel et al., 2015). Our behavioural findings showed that actions were estimated as occurring earlier when the Libet’s clock was displayed in near space as compared to far space. This influence of the clock’s proximity to the body on perceived action timing, despite no actual change in action initiation timing, replicates our previous findings (Kong et al., 2024). However, this spatial effect was absent in the perception of the tone’s timing in the Tone-only task, suggesting that the modulation of temporal perception is associated with voluntary action rather than altering time perception in general. This could suggest that spatial proximity enhances the saliency of actions, thus accelerating subjective time perception related to these actions, and also further prompts the question of how spatial proximity may shape the subjective experience of agency, and whether this modulation occurs through perceptual, motor, or decisional mechanisms. EMG results showed that spatial proximity increased the likelihood of earlier-initiated but longer-lasting (slower) movements (i.e., longer motor times) in a decision-reporting task without subsequent action outcome. As these results come from an exploratory analysis, their interpretation remains speculative at this stage. These results might indicate a generally earlier preparatory state under this task in the near space compared to the far space, aligning with the prior behavioral work of Blini and colleagues (2018), reinforcing the point that proximity to the body dynamically shapes sensorimotor engagement. Moreover, these results may reflect two complementary mechanisms: urgency and evidence accumulation (Steinemann et al., 2018). Earlier-initiated responses could arise from a heightened sense of urgency, related to both the proximity of the clock and the attentional focus on an earlier time point (i.e., the decision). In contrast, longer-lasting responses may reflect the greater need to accurately determine the timing of decision, which is inherently more demanding than reporting the timing of the action. Taken together, these findings emphasize the functional role of PPS not only in perception but also in voluntary action preparation and awareness. These results also highlight the necessity for further research investigating how proximity interacts with task demands, attentional states, and outcome expectations to modulate neural dynamics underlying voluntary action.

Note that we previously found that action initiation occurred earlier when the time marker (the clock) was displayed in near space (Kong et al., 2024). Here, we did not replicate this effect, likely due to methodological differences. Specifically, the present study employed longer intertrial intervals (ITI), more trials and a within-subject design, requiring participants to complete tasks in a randomized sequence across all tasks. This contrasts with a between-subject design of our earlier study, which featured shorter ITIs and fewer trials. These methodological changes likely contributed to a generally slower pace in the current study, potentially accounting for the discrepancy in action initiation timing. To investigate this further, we conducted a follow-up validation experiment with three conditions. The first condition replicated the Action-only condition from our previous study, with shorter ITIs (randomly chosen from 1s to 2s) and fewer trials (30 trials each for near and far). The second condition increased the ITI to match that of the current study (randomly chosen from 2s to 2.5s), while the third condition increased the number of trials to match that of the current study (50 trials each for near and far). Thus, the second and third conditions each differ from the first condition by only one parameter. The results showed that participants initiated the action significantly earlier in the near space compared to the far space in the first and third conditions. However, in the second condition, where only the ITI was increased, there was no significant difference in action initiation time between near and far spaces. Detailed results of this experiment are provided in the supplemental material (**Supp. Fig. S7**). These findings suggest that the increased ITI in the current study may have influenced movement times by reducing the urgency participants felt, thereby slowing overall response timing and eliminating the near–far difference in movement initiation. Importantly, the present results also indicate that PPS-related modulation of the RP does not necessarily translate into faster actions. Rather, consistent with recent evidence on the influence of awareness level on the RP (Gavenas et al., 2025; Parés-Pujolràs et al., 2023), PPS may influence the RP independently of motor urgency by modulating the awareness level during action preparation. Thus, PPS-related enhancement of RP activity may reflect increased access to—or integration of—awareness-related processes during motor preparation, rather than changes in behavioral motor urgency per se. Alternatively, it is also possible that PPS benefits from a privileged neural representation for both perception and action, independently of urgency or awareness.

The debate surrounding the RP centers on its interpretation: whether it directly indicates the preparation for volitional action, is an artifact of random neural fluctuations, or reflects higher order cognitive processes. Schurger et al. (2012) significantly contributed to this discussion by proposing that the RP represents a decision-making process where spontaneous fluctuations in neural activity accumulate until a threshold is reached, thereby triggering action execution. This model challenges the traditional view of the RP as a straightforward precursor to voluntary movement, instead suggesting that it reflects a stochastic accumulation process. Alongside this, Schultze-Kraft et al. (2021) showed that the RP cannot be suppressed, even when participants consciously attempt to do so via a neurofeedback system while still performing voluntary movements, suggesting an obligatory component to its generation. Moreover, empirical studies have shown that the RP is influenced by various factors (Maoz et al., 2019; Travers & Haggard, 2021). Our study aligns with this perspective, revealing that attentional focus and action outcomes along with spatial distance, specifically, the proximity of a clock, modulates the RP in task-specific ways. These findings underscore the RP’s sensitivity to situational variables, suggesting how certain contextual factors can influence neural excitability, bringing it to the action-triggering threshold, according Schurger et al.’s model, or modulating the temporal evolution and magnitude of motor preparation according to more traditional interpretations. Future studies should also investigate how RP effects relate to other neural markers of motor preparation, particularly beta-band activity over sensorimotor regions. Recent work has shown that the subjective feeling of readiness may link more closely with transient reductions in motor beta power than with RP amplitude itself (Gavenas et al., 2025; Parés-Pujolràs et al., 2023). Although this question lies beyond the scope of the present paper, our ongoing work specifically examines how beta activity and RP components reflect the dynamics of voluntary action preparation across different tasks.

One limitation of our study is the relatively low number of EEG electrodes, all positioned along the midline. Consequently, we may not have captured the maximal RP amplitude, which can vary across individuals and may peak at more frontal sites such as FCz under certain conditions (Travers & Haggard, 2021). Therefore, we cannot rule out the possibility that some of the observed modulation in RP magnitude partly reflects changes in RP topography across experimental conditions. Nevertheless, the present results clearly show that the experimental manipulations affected the RP waveform, whether it primarily impacts the temporal dynamics of its buildup at a specific location or across its spatial distribution.

Overall, this study discloses that attentional focus, action outcomes and spatial proximity each play a distinct and an interacting role in shaping RP and action awareness. The RP emerges as a highly flexible neural marker, adapting to task demands, internal higher cognitive processes and external environmental context, emphasizing how attentional focus on intention to act modulates early RP—traditionally considered an unconscious stage of neural readiness. Future research should further explore these dynamics across various voluntary action contexts, and how these factors interact with sense of agency and action decision-making processes.

## Author Contributions

G.K. and A.F. led the conceptualization of the study. The study design was led by G.K., with supporting contributions from M.V. and A.F. G.K. took the lead in preparing the experimental setup and software, supported by B.B., C.D., and M.V. Data collection was conducted by G.K. and B.B. Data analysis was led by G.K., with support from B.B., A.F., and M.V. Data interpretation was led by G.K., M.V. and A.F. The original draft was written by G.K. Revisions were led by G.K., M.V., and A.F., with supporting input from L.M. and F.P. All authors reviewed and approved the final version of the manuscript.

## Acknowledgements

The authors would like to thank the editor and the reviewers for their insightful and constructive feedback; Cheryne Aberkane for the help with pilot data collection; Cécile Fabio for the support with EEG techniques; Mélanie Ducellier for the validation data collection; Sonia Alouche, Jean-louis Borach for administrative support.

## Ethics

This research complies with the Declaration of Helsinki (2023), aside from the requirement to preregister human subjects research, and received approval from a local ethics board (n°21-772, IRB00003888, IORG0003254, FWA00005831).

## Data accessibility

All primary data, analysis scripts and materials are publicly available on the Open Science Framework repository (https://osf.io/tjus4/).

