## Supplemental material for "Spatial distance and temporal attentional focus modulate voluntary action preparation and awareness"

**This includes:**

Figure S1. Illustration of the EMG onset detection method.

Figure S2. Computer-registered keypress movement initiation times for all tasks involving keypresses.

Figure S3. Movement initiation time distributions by distance.

Figure S4. Movement initiation time distributions by task.

Figure S5. Histograms of motor times across tasks.

Figure S6. The time-course of RP across different experimental tasks with epochs aligned on EMG-detected movement onset and without baseline correction.

Figure S7. Validation experiment for movement initiation time.

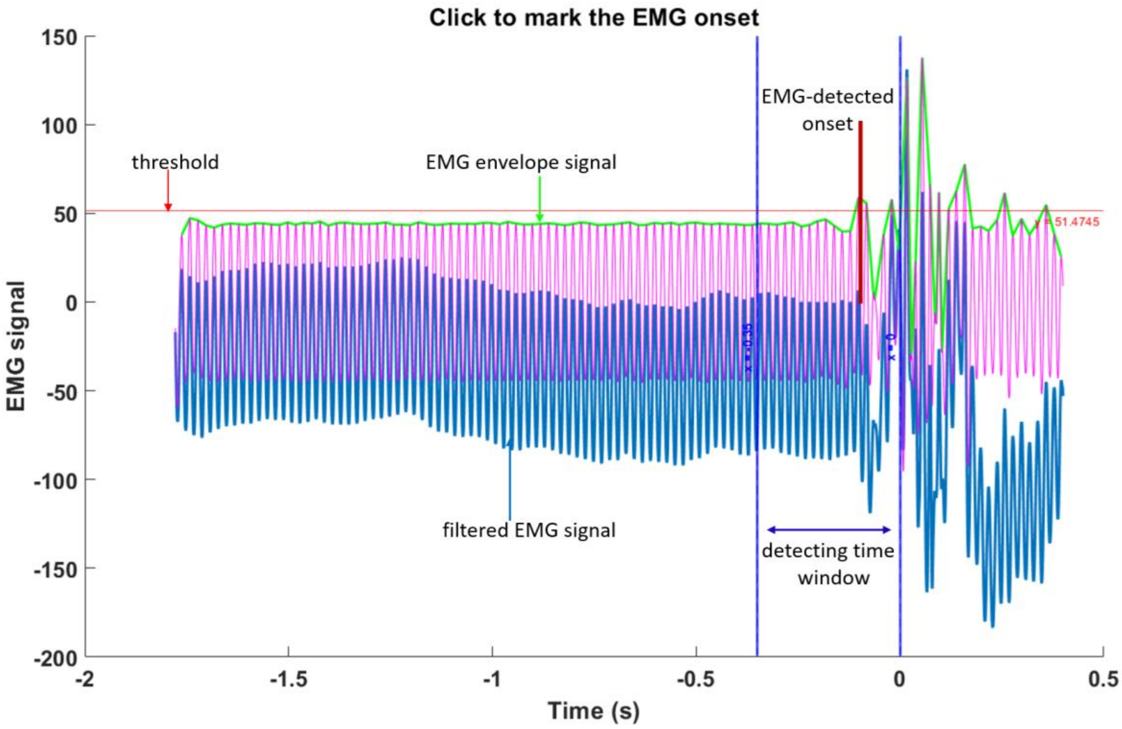


**Figure S1. Illustration of the EMG onset detection method.** The blue waveform represents the original EMG signal and the purple waveform the filtered EMG signal, processed to remove noise and enhance signal clarity. The EMG envelope signal (green line) is obtained by rectifying and smoothing the filtered signal, providing a more stable representation of muscle activation levels. A threshold (red horizontal line, 50% of the Peak-Baseline difference) is used to determine the onset of muscle activity. The detection period is marked by two vertical blue dashed lines at x=−0.35 and x=0 (computer-registered keypress onset), defining the time window in which EMG onset is assessed. The EMG-detected movement onset is marked by the vertical red line. If the automatically detected onset is not optimal, the user is prompted to manually select the onset point within this window for further analysis.

### Movement initiation time

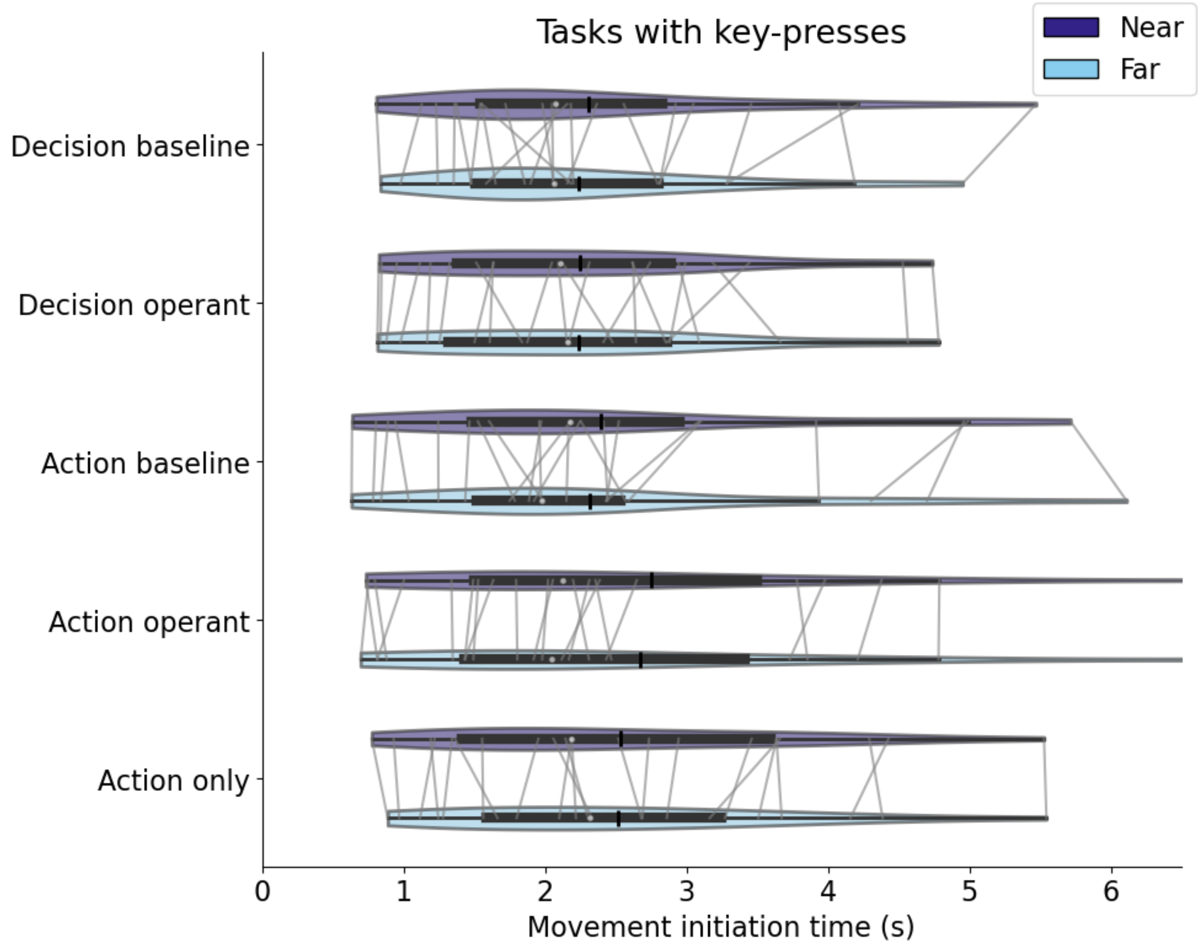


**Figure S2. Computer-registered keypress movement initiation times for all tasks involving keypresses.** The light blue and purple violins distinguish between near and far distances, respectively. Each violin's width reflects the probability density of data points across different movement initiation time values on the x-axis, offering a visual portrayal of data distribution within each task. The white dot within each violin represents the median movement initiation time, the thick horizontal line represents the interquartile range, and the vertical black bar represents the mean movement initiation time within each task.

For the computer-registered keypress movement initiation time (**Fig. S2**), the paired t-test for the Action-only task showed no significant difference between the clock presented near (Mean ± SE = 2.51 ± 0.24 s) and far (Mean ± SE = 2.48 ± 0.23 s, t (24) = .65, p = .52). The 2 (report instruction: decision vs. action) × 2 (distance: near vs. far) × 2 (tone presence: baseline vs. operant task) three-way mixed design ANOVA showed no significant main effect of spatial distance (F (1,24) = 0.25, p = 0.62, $\eta_p^2$ = .01), of report instruction (F (1,21) = 0.72, p = .41, $\eta_p^2$ = .03), and of tone presence (F (1,24) = 0.52, p = .48, $\eta_p^2$ = .02) and no interaction was significant (p > 0.41). These results suggest that PPS did not significantly influence the overall timing of keypresses in all tasks. Similarly, results from EMG-detected movement onset times also showed no significant influence of PPS on movement onset timing in all tasks.

### Movement initiation time distributions by distance

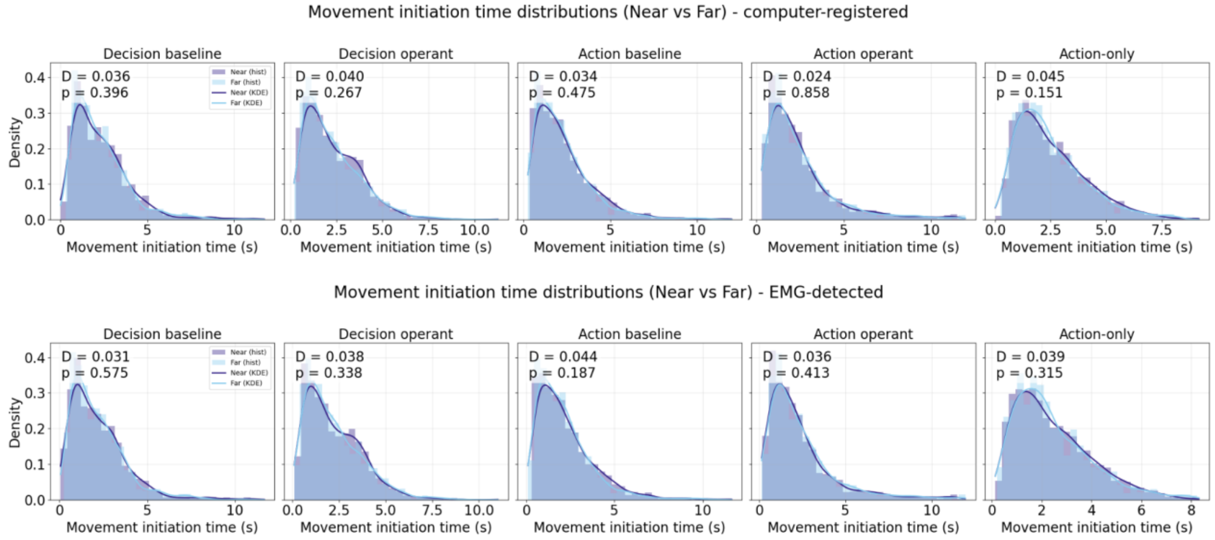


**Figure S3. Movement initiation time distributions by distance.** Distributions of movement initiation times for Near (purple) and Far (light blue) conditions for the five tasks: Decision baseline, Decision operant, Action baseline, Action operant, and Action-only. Panels show probability density estimates (kernel density functions, KDEs) overlaid on histograms of individual movement initiation times (in seconds), pooled across participants and trials. The top panel displays computer-registered movement initiation times, and the bottom panel shows EMG-detected movement initiation times. The Kolmogorov–Smirnov statistics (D, p) reported in each panel indicate that no significant differences were found between Near and Far conditions in any task (all p > 0.15).

### Movement initiation time distributions by task

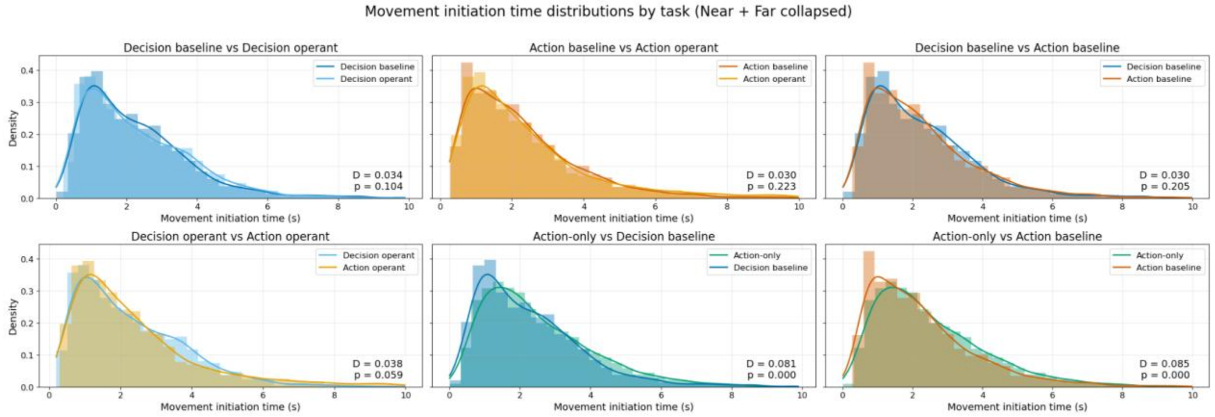


**Figure S4. Movement initiation time distributions by task.** Each subplot compares the movement initiation time (MIT) distributions between two task conditions using density histograms and KDEs. The top row shows comparisons between baseline and operant versions of the Decision task (left), the Action task (middle), and Decision vs. Action baseline tasks (right). The bottom row shows comparisons between Decision vs. Action operant tasks (left), Action-only vs. Decision baseline (middle), and Action-only vs. Action baseline (right). For each comparison, the Kolmogorov–Smirnov statistics (D, p) are reported to quantify whether the two distributions differ significantly. Distributions largely overlap across Decision and Action tasks, with no significant differences except when the Action-only condition is involved, which exhibits a distinct distribution characterized by slower and less sharply peaked initiation times.

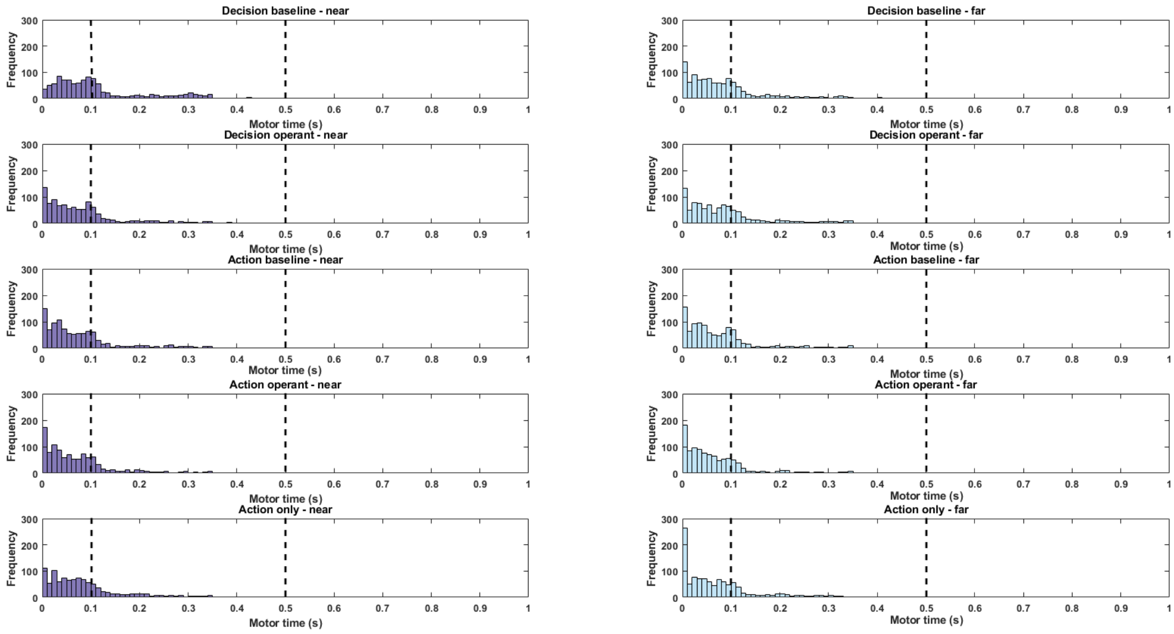


**Figure S5. Histograms of motor times (i.e., keypress-EMG movement initiation time (MIT) differences) across tasks.** The distribution of motor times within each task: Decision Baseline, Decision Operant, Action Baseline, Action Operant, and Action-only, with both near and far distance settings. The x-axis represents the motor time in seconds, while the y-axis represents the frequency of occurrences. The histograms show that most motor times fall within the critical range of 0 to 0.1 seconds, but an interesting second part of the distribution occurred within 0.1-0.5 s. The dashed vertical lines highlighted this critical range (0.1 to 0.5 seconds) used for further analyses (see “Movement initiation times and motor times” in the Results of the main manuscript).

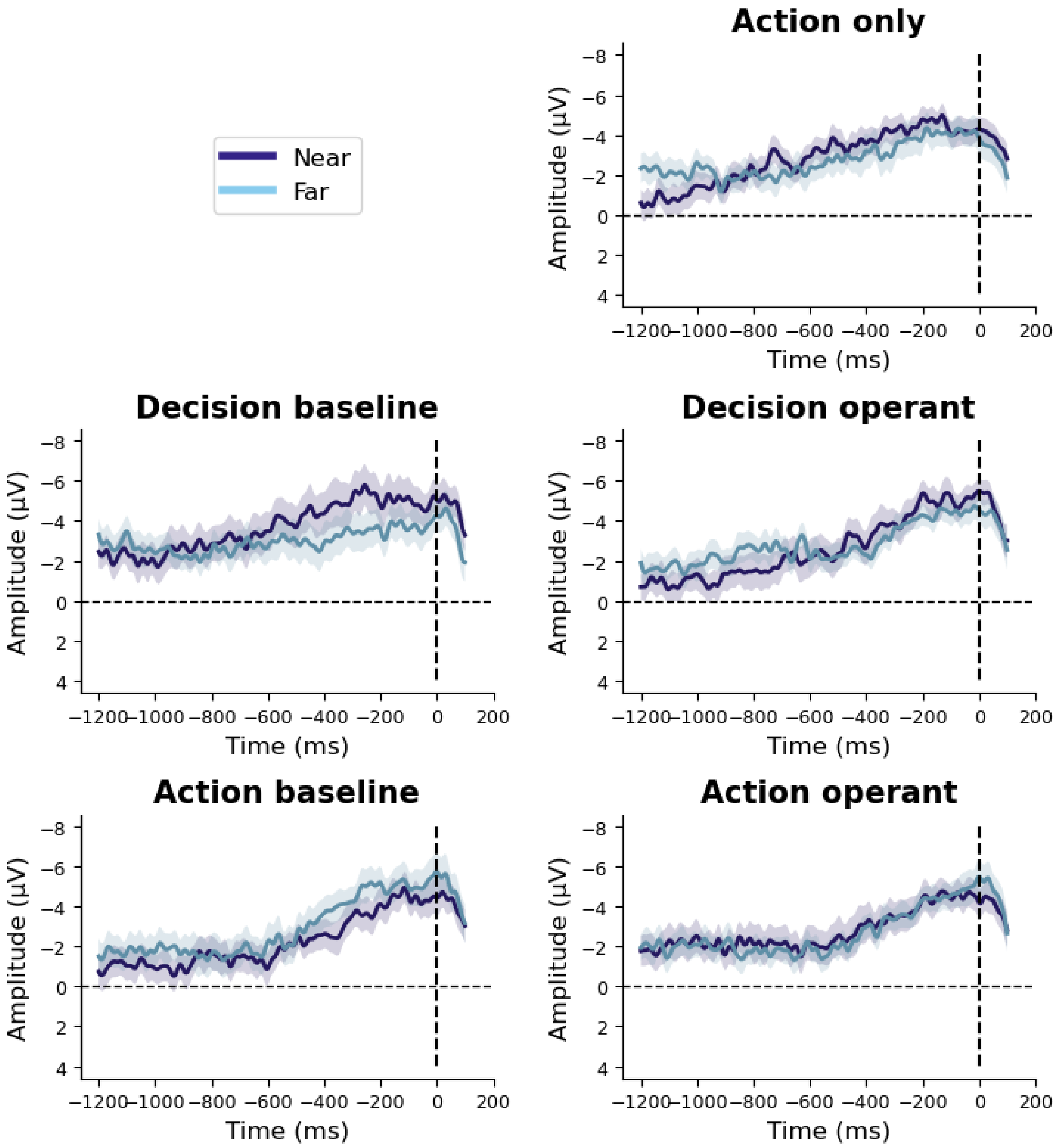


**Figure S6. The time-course of RP across different experimental tasks with epochs aligned on EMG-detected movement onset and without baseline correction.** The vertical dashed line at time zero represents the EMG-detected movement onset. The light blue line indicates the RP when the clock was in close proximity to the participants, while the purple line depicts the RP when the clock was positioned further away. The shaded regions surrounding each line represent standard deviation errors of the mean.

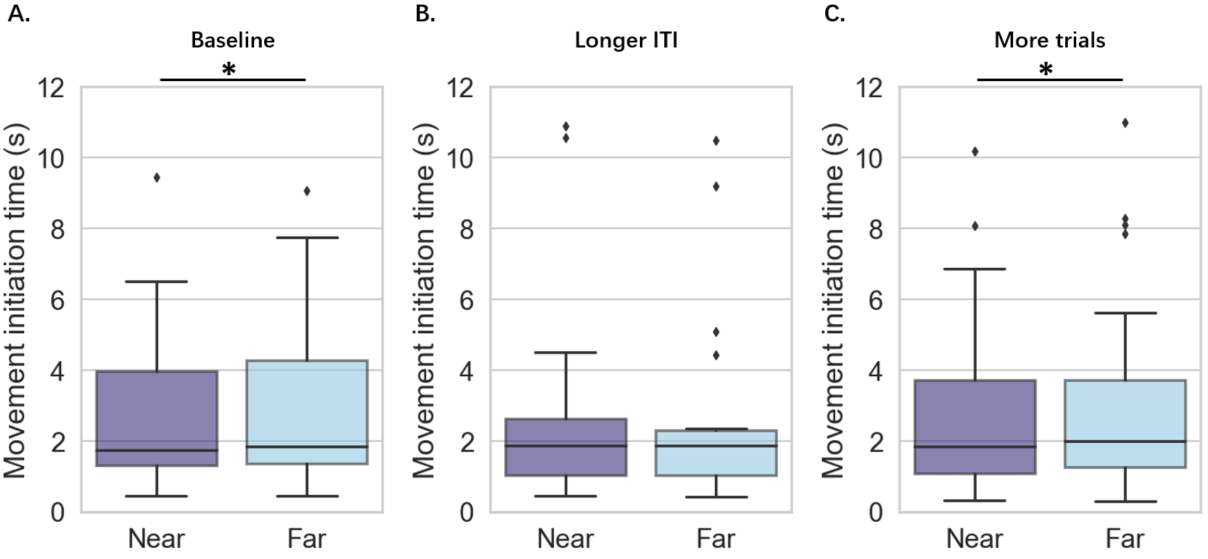


**Figure S7. Validation experiment for movement initiation time.** To determine whether and which factors—longer inter-trial intervals (ITI, **Fig. S7B**) or a higher number of repeated trials (**Fig. S7C**)—affect movement time, we conducted a follow-up validation experiment with three conditions from 20 participants (12 females, mean age = 27 years, range 20–47 years). The first condition (**Fig. S7A**) replicated the Action-only condition from our previous study (Kong et al., 2024), with shorter ITIs (randomly chosen from 1s to 2s) and fewer trials (30 trials each for near and far). The second condition (**Fig. S7B**) increased the ITI to match that of the current study (randomly chosen from 2s to 2.5s), while the third condition (**Fig. S7C**) increased the number of trials to match that of the current study (50 trials each for near and far). Thus, when comparing the first and second conditions, only one parameter differed between them. The results showed that participants initiated the action significantly earlier in the near space compared to the far space in the first and third conditions ($t(19) = -2.24$, $p = 0.037$; $t(19) = -2.34$, $p = 0.03$, respectively). However, in the second condition, where only the ITI was increased, there was no significant difference in movement time between near and far spaces ($t(19) = 1.26$, $p = 0.22$). These findings suggest that the increased ITI in the current study may have influenced movement times by reducing the urgency participants felt, thereby slowing down the overall process. This likely explains the discrepancy in movement times observed between our previous study and the current one.
